# An amygdala-cortical circuit for encoding generalized fear memories

**DOI:** 10.1101/2025.01.15.633190

**Authors:** Carly J. Vincent, Ricardo Aguilar-Alvarez, Samantha O. Vanderhoof, David D. Mott, Aaron M. Jasnow

## Abstract

Generalized learning is a fundamental process observed across species, contexts, and sensory modalities that enables animals to use past experiences to adapt to changing conditions. Evidence suggests that the prefrontal cortex (PFC) extracts general features of an experience that can be used across multiple situations. The anterior cingulate cortex (ACC), a region of the PFC, is implicated in generalized fear responses to novel contexts. However, the ACC’s role in encoding contextual information is poorly understood, especially under increased threat intensity that promotes generalization. Here, we show that synaptic plasticity within the ACC and signaling from amygdala inputs during fear learning are necessary for generalized fear responses to novel encountered contexts. The ACC did not encode specific fear to the training context, suggesting this region extracts general features of a threatening experience rather than specific contextual information. Together with our previous work, our results demonstrate that generalized learning about threatening contexts is encoded, in part, within an ascending amygdala-cortical circuit, whereas descending ACC projections to the amygdala drive generalized fear responses during exposure to novel contexts. Our results further demonstrate that schematic learning can occur in the PFC after single-trial learning, a process typically attributed to learning over many repeated learning episodes.

## Introduction

To survive, animals must use experience to adapt flexibly to changing and uncertain conditions. Animals typically use generalization as an adaptive process to assess if new objects or situations are likely to produce the same outcome as those previously experienced. Thus, unlike a lack of discrimination, which is the failure to detect a difference between two stimuli, generalization is an active cognitive process that promotes appropriate behavioral action across contexts and stimuli (Shepard, 1987). Likewise, generalizing fear responses is evolutionarily adaptive because it supports survival by enabling animals to avoid threatening situations similar to ones experienced previously but come at the cost of missing out on food resources or access to mates. In fear conditioning protocols, generalization occurs when animals, including humans, produce fear responses to contexts or cues that are similar but distinctly different than those previously associated with an aversive stimulus; this is the basis for the overgeneralization observed with anxiety and stress-related disorders [e.g., post-traumatic stress disorder (PTSD)] (Lissek et al., 2008; 2010). Indeed, fear generalization is recognized as a transdiagnostic mechanism for anxiety disorders (Cooper et al., 2022). Studies report that increasing threat intensity (e.g., shock) results in more generalization in rodents and humans (Ortiz et al., 2019; Laxmi, et al., 2003; Dunsmoor et al., 2017), but how fear spreads across multiple stimuli during increased threat is not well-established (Ghosh et al., 2014). Furthermore, whereas valuable mechanisms related to encoding and expressing fear discrimination to extensively trained discriminatory contexts and tones (i.e., CS^+^ and CS^-^) have been identified (e.g., Ghosh et al., 2014; Martin-Fernandez et a., 2023; Stujenski et al., 2022; Besnard et al., 2019; 2020; Likhtik et al., 2014), how rapid encoding for generalizing to never-experienced stimuli or circumstances after single-trial learning are not well defined.

The medial prefrontal cortex (mPFC) extracts general features of an experience rather than detailed contextual or spatial information (Yu et al., 2018; Martin-Fernandez et al., 2023). The anterior cingulate cortex (ACC, Brodmann’s area 24A, 24B), a subregion of the mPFC, is involved in the expression of generalized fear memories (Cullen, et al., 2015; Adkins, et al., 2019; Einarisson, et al., 2015; Ortiz, et al, 2019) and storing remote memories (Einarisson et al., 2012; Frankland et al., 2004; Restivo et al., 2009), but its role in encoding context representations is less clear. The nucleus reuniens (RE) is a thalamic structure that connects the PFC to the hippocampus and is important in acquiring hippocampal-dependent contextual memories (Xu and Sudhof, 2013; Ramanathan et al., 2018). Without a functioning RE, acquired memories are hippocampal-independent and presumably rely on cortical regions to be encoded (Ramanathan et al., 2018). However, even with a fully functioning RE, the mPFC supports generalized representations, suggesting it may operate in parallel with an RE-hippocampal network to encode different task features (Yu et al., 2018; Martin-Fernandez et al., 2023). The basolateral amygdala (BLA) sends prominent projections to the mPFC (Hoover and Vertes, 2007; McGarry and Carter, 2016), but how these inputs to the ACC are involved in encoding context and threat information is poorly understood, especially under conditions that promote generalized memories (Ortiz et al., 2019; Laxmi et al., 2003; Dunsmoor et al., 2017). To examine these questions, we used pharmacological and chemogenetic-mediated circuit manipulations to uncover the role of the ACC in encoding highly salient experiences that generalize across contexts.

## Methods & Materials

### See the supplement for detailed methods

#### Animals

Experiments used 7-12 week, male and female F1 129B6 hybrids derived from crossing C57BL/6J males with 129S1/SvImJ females. All mice were generated in a breeding colony at the University of South Carolina School of Medicine. All mice were housed in groups of two-five per cage, had ad libitum access to food and water, and maintained on a 12:12 light-dark cycle. All procedures were conducted in a facility accredited by the American Association for Laboratory Animal Care (AALAC), per the National Institutes of Health guidelines, and with approval by the University of South Carolina Institutional Animal Care and Use Committee (IACUC) guidelines.

#### Surgeries

##### Cannulations

Mice were given a subcutaneous injection of Rymadil (2.5mg/kg) prior to anesthesia. Following drug administration, mice were anesthetized with inhaled isoflurane (3% induction: 1-1.5% maintenance). Mice were then tested for lack of motor responses before placement into the stereotaxic apparatus (David Kopf Instruments, Tujunga, CA). An incision on the scalp was made and measurements of lambda and bregma landmarks were determined. For the pharmacology experiments, a unilateral burr hole was drilled using the following coordinates for the ACC: (+.80mm AP, +.70mm ML, -1.75mm DV). A 1mm projection needle inserted into a 4mm cannula (Plastics One, Inc.) set at a 14° angle was guided to the appropriate position in relation to bregma. For the circuit activation experiment, mice were cannulated over the Basolateral Amygdala (BLA): (-1.6mm AP, +3.4mm ML, -4.9mm DV). A single 3mm projection needle inserted into a 4mm cannula was used. Following surgery, mice were removed from the apparatus, placed into a clean cage, and monitored until fully awake.

##### Virus Infusions

All mice were given a subcutaneous injection of Rymadil (2.5mg/kg) prior to anesthesia. Following drug administration, mice were anesthetized with inhaled isoflurane (3% induction: 1-1.5% maintenance). Mice were then tested for lack of motor responses prior to placement into the stereotaxic apparatus (David Kopf Instruments, 129 Tujunga, CA). An incision on the scalp was made and measurements of lambda and bregma landmarks were determined. Bilateral burr holes were drilled using the following coordinates for the Prelimbic Cortex (PL): (+ 1.90mm AP, +.60mm ML, -2.25mm DV) at a 12 ° angle; Anterior Cingulate Cortex (ACC): (+.80mm AP, +.70mm ML, -1.75mm DV) at a 14 ° angle; BLA (-1.6mm AP, +3.4mm ML, -4.9mm DV). For PL inactivation, pAAV-hSyn-hM4D(Gi)-mCherry (AAV8) (Addgene) or pAAV-hSyn-EGFP (AAV8) (Addgene) were bilaterally infused into the PL at a volume of 0.2µl per side, at a rate of 0.1µl per minute. For BLA-ACC circuit inactivation, a retrograde pAAV-Ef1a -cre (AAVrg) (Addgene) was bilaterally infused into the ACC at a volume of 0.6µl per side at a rate of 0.2µl per minute (Ramsaran, et al., 2023; Kol, et al., 2020; Penzo, et al., 2015). Following the ACC infusion, pAAV-hSyn-DIO-hM4D(Gi)-mCherry (AAV8) or pAAV-hSyn-DIO-mCherry (AAV8) (Addgene) was bilaterally infused into the BLA at a volume of 1.0µl per side at a rate of 0.2µl per minute. For the BLA-ACC circuit activation experiment, pAAV-hSyn-hM3Dq-mCherry (AAVrg) or pAAV-CaMKIIa-EGFP (AAVrg) was infused into the ACC at a volume of 0.2µl per side at a rate of 0.1µl per minute (Ortiz, et al., 2019). Five weeks later mice were cannulated over the BLA and allowed 1 week to recover prior to behavioral testing. This enablded inactivation of BLA terminals in the ACC as we have done previously (Ortiz et al., 2019).

#### Behavioral Procedures

##### Context Fear Conditioning

Mice were trained using contextual fear conditioning in four identical conditioning chambers (12” W x 12” D x 12”H) containing two Plexiglas walls, two aluminum sidewalls, and a stainless-steel grid-shock floor (Colbourn Instruments, Allentown, PA). The training context consists of the conditioning chamber with a polka-dot insert attached to the rear Plexiglass wall, white noise (70db), dim illumination, and the stainless-steel grid floors cleaned with 70% ethanol. Mice received two days of pre-exposure for 5 minutes. Then, the following day, the mice underwent nine minutes of fear conditioning by pairing the training context with a series of 5 un-signaled foot shocks (1.0 mA) separated by 90-second inter-shock intervals (ISI). Measures of freezing were recorded using the Freezeframe5 (Actimetrics). Mice were then tested for fear expression by examining freezing across the testing period consisting of a 5-minute exposure to the training context and a novel context in a counterbalanced design with 72 hours between tests and were matched based on the last 30 seconds of freezing during training. We have previously demonstrated that this procedure produces no order effects (Ortiz et al., 2019). The novel context had no background element, a flat floor, a fan (60db), IR lights and was cleaned with 2% quatricide (Pharmacal). For experiments that assessed activity-regulated cytoskeletal-associated protein (Arc) expression in the ACC, one group of mice was exposed to an immediate shock procedure to take advantage of the immediate shock deficit (Wiltgen, et al., 2001; Fanselow, 1990). When rodents are shocked immediately after being placed into a conditioning context, they do not exhibit freezing upon re-exposure, presumably because they have not had enough time to develop a representation of the conditioning context (Rudy & O’Reilly, 2001). Before immediate shock, mice received two days of pre-exposure, each for five minutes, to a novel cage to control for pre-exposure in the group of mice receiving standard context fear conditioning. Then, the following day, mice underwent the immediate shock procedure by being placed in the training context and were administered 5 un-signaled foot shocks in 10 seconds (1.0 mA). After the last shock, mice were immediately removed from the chamber and returned to their cage.

##### Explicitly Unpaired Context Fear Conditioning and Immediate Shock Procedure

To demonstrate the involvement of associative learning in generalization, we utilized an explicitly unpaired context fear conditioning procedure and, separately, an immediate shock procedure. For explicitly unpaired context fear learning, the mice were fear-conditioned in an alternative conditioning chamber and then exposed to our standard training context six hours later to serve as an ‘explicitly unpaired’ context. Mice were tested for fear in the novel and standard training contexts as above. The alternative training context consisted of operant conditioning chambers (Med-Associates; 8.25” W x 7.5” D x 11” H), with a BLAck and white striped insert on the rear wall, a vanilla-scented cotton ball placed in the bottom tray of the chamber, and stainless-steel grid floors that were cleaned with 70% ethanol prior to and in between conditioning. Mice were conditioned in the dark. The training conditions were the same as above. Mice were exposed to the standard training context for 9 minutes, the same length as the fear-conditioning procedure. The following day, mice were tested in the standard training context or a novel context (same as above) for 5 minutes and were matched based on the last 30 seconds of freezing during context fear conditioning. Mice underwent testing 72 hours after the first test in the opposite context in which they were originally tested. Additionally, to assess fear to the alternative training context (operant chambers) mice underwent a 5-minute test 6 days following the last exposure. Freezing was recorded using the Freezeframe5 (Actimetrics), or ANY-Maze 4.99 software (Stoelting, Wood Dale, IL).

When assessing the need for a context representation to promote generalization, mice were run through an immediate shock procedure. Mice received two days of handling. The following day animals were placed in the standard training context as previously described and received 5-unsignaled foot shocks (1.0mA) over 10 seconds. Twenty-four hours later, mice were returned to the training context and received a 9-minute context exposure. The following day, mice were tested for fear expression in the novel context (elements previously described).

##### Open Field Test

For the assessment of BLA-to-ACC circuit inactivation on locomotor behavior, mice were placed in four identical 45cm X 45cm white acrylic open-field arenas. Lighting was adjusted to 10-12 lux. Each arena was cleaned with 10% ethanol prior to and in between uses. Mice were recorded for 10 minutes, and distance traveled was assessed using Ethovision15 software.

#### Drug Preparation

Lidocaine HCL (Sigma) was dissolved in phosphate-buffered saline at a concentration of 4% w/v and adjusted to a pH of 7.0 (Frankland, et al., 2004; Cullen, et al., 2015). DL-AP5 (HelloBio) was dissolved in 0.9% saline at a concentration of 45mM (2.7 μg/side) (Jasnow, et al., 2004; Walker, et al., 2005; Adkins, et al., 2019). Clozapine-N-oxide dihydrochloride (CNO) (HelloBio) was dissolved in 0.9% saline at a dose of 5mg/kg, for intraperitoneal injections, or 0.9% saline at a concentration of 650 μM for intracranial infusions (Ortiz, et al., 2019).

#### Intracranial Infusions

For all pharmacological experiments, mice received infusions immediately before or after contextual fear conditioning. Mice were transported from the animal colony to a room adjacent to the fear conditioning room, where infusions occurred. For post-training infusions, mice were removed from the fear conditioning chambers, placed into their home cage, and transported to the infusion area. For intracranial infusions, mice were placed in a clean cage, and a 33-gauge internal infusion needle attached to polyethylene tubing (PE-20) was inserted into the guide cannula. A 5μL Hamilton syringe attached to a micro-infusion pump from Harvard Apparatus was used to infuse the drug or vehicle solution. All mice received a 0.2μL infusion of vehicle or drug unilaterally into the Anterior Cingulate Cortex at a rate of 0.1μL/min. For BLA infusions, mice received 0.2μL (per side) of CNO bilaterally into the BLA at a rate of 0.1μL/min. After the infusions, needles were left in place for 2 minutes to allow for diffusion of the drug. Afterward, dummy cannulas were inserted, and mice were returned to their home cage or conditioning chambers.

#### Cannulation Site Verification

After behavioral testing, mice were infused with 4% fluro-gold (Flurochrome, LLC) at 0.05μL/minute for a total volume of 0.1μL via guide cannula. Brains were extracted and stored in a -80°C freezer until slicing. Sections were sliced at 40μm on a cryostat (Microm HM 500) and placed onto superfrost + slides. Sections were cover-slipped with prolonged diamond mounting media and sites were examined using an epi-fluorescent microscope (Leica). Mice whose sites were outside of the ACC or BLA were excluded from statistical analyses.

#### Immunohistochemistry

To verify virus expression after behavioral testing, mice were deeply anesthetized and transcardially perfused using 0.9% saline, followed by 4% paraformaldehyde. Brains were post-fixed for 24 hours, then transferred to 30% sucrose solution until fully submerged. Brains were sliced on a freezing microtome (Leica) at 40μM thickness and stored in an ethylene glycol/sucrose antifreeze solution at -20°C until processing. Sections were then stained for mCherry using immunohistochemistry. All sections were incubated in hydrogen peroxide (0.08% and 0.3%), washed in PBS, and then incubated in rabbit anti-mCherry primary antibody (1:30,000; abcam, ab167453) for 1 hour at room temperature then 4° C for 48 hours with constant agitation. After primary antibody incubation, tissue was washed in PBS, then incubated with a goat anti-rabbit biotinylated secondary antibody (1:500; Jackson Immuno, 111-005-003). After, avidin-biotin complex (ABC; Vector Laboratories) was applied to the tissue for 1 hour. Following ABC application, tissue was washed briefly in PBS and sodium acetate (0.175 M) and visualized by adding Ni-enhanced 3, 3’-Diaminobenzidine (DAB). Ni-DAB solution was applied for 20 minutes. Following Ni-DAB enhancement, tissue was washed in sodium acetate (0.175 M and PBS. Afterward, the tissue was mounted and dehydrated via increasing changes of ethanol and application of xylenes for clearing. Sections were coverslipped using DPX mountant and visualized on a light microscope. Sections that included hM4Di-mCherry expression outside of the BLA or lacked fiber staining in the ACC were excluded.

For immunohistochemical detection of activity-regulated cytoskeletal-associated protein (Arc), mice were perfused 60 minutes after context fear conditioning. Brains were post-fixed for 24 hours, then transferred to 30% sucrose solution until fully submerged. Brains were sliced on a freezing microtome (Leica) at 40μM thickness and stored in an ethylene glycol/sucrose antifreeze solution at -20°C until processing. Sections were removed from cryoprotectant solution and washed in PBS. Following washes, sections were incubated in rabbit anti-Arc primary antibody (1:30,000; abcam, ab183183) for 1 hour at room temperature, then 4° for 48 hours with constant agitation. Following incubation, sections were washed with PBS and incubated in a goat anti-rabbit biotinylated secondary antibody (1:500; Jackson Immuno, 111-005-003) for 1 hour at room temperature. Following incubation, sections were washed and placed in Avidin-Biotin Complex (ABC), and washed and enhanced with Ni-DAB, as described above. Afterward, tissue was mounted and dehydrated via increasing changes of ethanol and application of xylenes for clearing. Sections were then coverslipped using DPX mountant, then visualized on a light microscope.

#### Analysis of Arc Expression

To assess Arc protein expression in the ACC, images were taken on a light microscope (Leica DM 500B) using a 20X objective. Images were analyzed using FIJI (NIH Image) to set a region of interest (ROI) for analysis. The ROI was 200μm x 200μm section set in the ACC. Landmarks were used to guide consistency in placement across slices (Pollack, et al., 2018) per the mouse brain atlas (Paxinos and Watson, 2019). Four non-consecutive hemispheres were analyzed by manually counting cells within the ROI (Pollack, et al., 2018). Researchers were blinded to treatment conditions throughout the counting process.

#### Statistical Analyses

All data were analyzed using GraphPad Prism statistical software (GraphPad 10) using unpaired t-tests, between subjects, or repeated measures two-way Analysis of Variance (ANOVA). Statistically significant ANOVAs were followed with Tukey’s or Sidak’s post-hoc analyses. Please refer to Tables 1 and 2 in the supplemental materials for statistical details.

## Results

### Plasticity in the anterior cingulate cortex is engaged during strong context fear conditioning

To investigate the role of the ACC during learning in encoding generalization, we first used activity-regulated cytoskeletal-associated protein (Arc) expression after different fear conditioning parameters as an indicator of neuroplasticity. Arc is a protein involved in learning and memory and is a reliable indication of neuroplasticity (Vazdarjanova et al., 2006; Carpenter-Hyland et al., 2010). Mice were trained using five un-signaled foot shocks in a contextual fear conditioning procedure, an immediate shock procedure to control for shock experience, or a home cage control procedure. During contextual fear conditioning, mice increased post-shock freezing with each shock presentation [F (2.204, 17.63) = 61.68, p < 0.0001] (Figure 1B). Arc staining in the ACC was significantly greater following 5 shock context fear training compared to immediate shock and home-cage control groups [F (2, 17) = 53.62, p < 0.0001] (Figure 1C). Next, we conducted an explicitly unpaired context fear conditioning experiment to demonstrate that fear generalization is not solely reliant on the strength of the shock alone but also depends on associative learning between the shock and the similarities of the contextual elements that are presented during training and testing (Thomas, 1981; *for review see*, Riccio, et al., 1984). Mice were trained in the context fear conditioning procedure described above but in an alternative training context with different cues (Med Associates 8.25” W x 7.5” D x 11” H) and no illumination. Six hours after training, they were placed into the standard training context, which served as an ‘explicitly unpaired’ context for nine minutes. The average freezing was significantly higher in the ‘alternative training’ context compared to the ‘explicitly unpaired’ context, as expected [t (10) = 10.56, p < 0.0001] (Figure 2B). Mice were tested for fear expression in the ‘explicitly unpaired’ context and a ‘novel’ context in a counterbalanced design. Six days later, mice were returned to the ‘alternative training’ context to assess freezing to the context in which they were trained (Figure 2C). There was a significant effect of context [F (1.915, 16.28) = 27.70, p < 0.0001]; during fear expression tests, mice displayed significantly higher freezing in the ‘alternative training’ context compared to the ‘explicitly unpaired’ and ‘novel’ contexts (Tukey’s MC test p = 0.0132; p < 0.0001). To demonstrate that shock exposure alone does not produce generalization and requires some representation of the training context, we utilized an immediate shock procedure. Mice were exposed to the immediate shock procedure as previously described. Twenty-four hours later mice were returned to the training context where they received a 9-minute context exposure. The next day they were tested for fear expression to the novel context. All mice showed low freezing to the novel context, indicating no context generalization (Figure 2C). Overall, these experiments demonstrate that increased shock intensity alone during strong training is not sufficient to produce generalization and suggest that associative learning between the shock and the similarities of the contextual elements present during training and testing is necessary for generalized learning.

**Figure 1:**
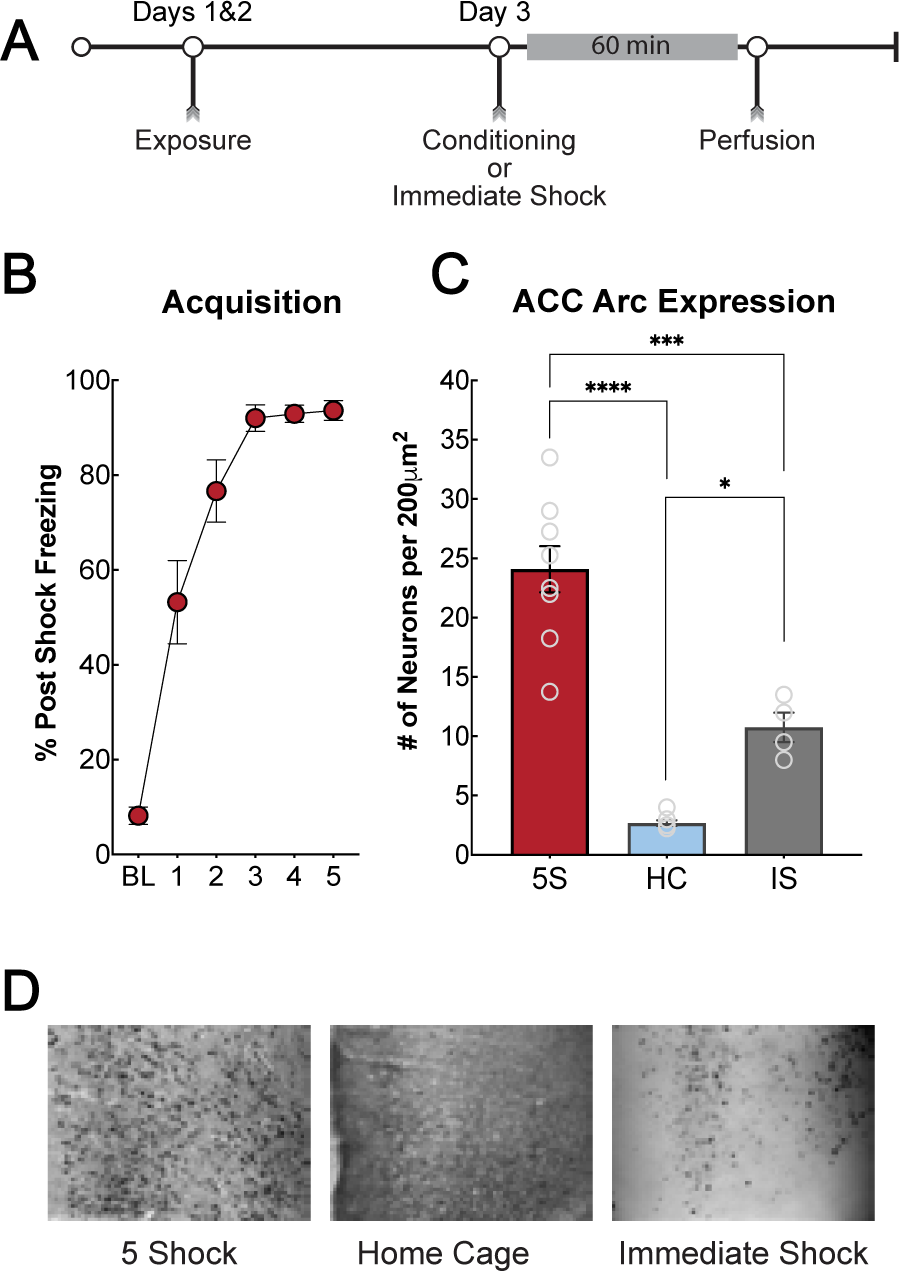
Arc expression in the ACC is increased during strong context fear conditioning. **A**: Timeline for behavior and Arc immunohistochemistry experiment. Mice underwent contextual fear conditioning as described. Sixty minutes after training, mice were perfused, and brain tissue was processed for immunohistochemistry to assess the expression of the plasticity-associated protein Arc. **B:** Post-shock freezing during acquisition increased after successive presentations of the 5 un-signaled foot shocks. **C:** Quantification of Arc expression in the ACC after strong training, home cage, or immediate shock with the same parameters as strong training (5 shocks, 1 mA). Arc expression was significantly greater in the ACC in mice trained with 5 unsignaled foot shocks (5S) compared to home cage controls (HC) and immediate shock controls (IS). **D:** Representative images of Arc immunohistochemistry in the ACC. From left to right, a representative image of the 5-shock training, home-cage controls, and immediate shock controls.

**Figure 2:**
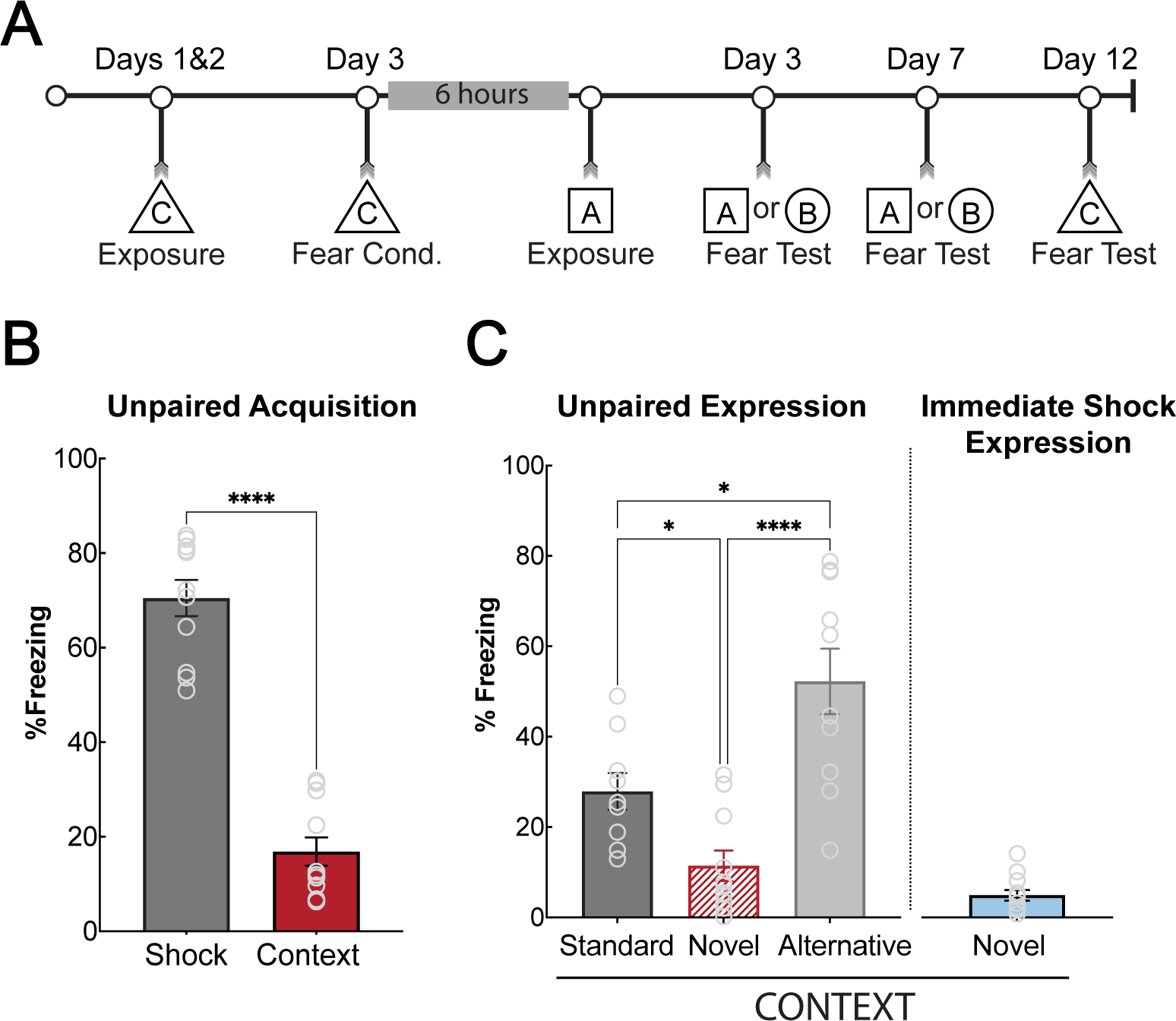
Generalization requires associating a general context representation and threat. **A:** Timeline for the unpaired context fear conditioning experiment and the immediate shock procedure. Mice were fear conditioned in an ‘alternative’ training context (triangle, C), then exposed to the ‘standard’ training context (square, A) six hours later. Mice were then tested for fear in a counterbalanced design in the ‘standard’ training and novel contexts (circle, B). They were then tested in the ‘alternative’ training context (C). For the immediate shock procedure, mice were placed in the training context and immediately received 5 footshocks and were immediately removed from the context. Twenty-four hours later, mice were exposed to the training context for nine minutes and one day later were tested for fear generalization in a novel context. **B:** Acquisition for unpaired context fear conditioning. Mice display higher freezing in the ‘alternative’ training context during training compared to the ‘standard’ training context exposure [t (10) = 10.56, p < 0.0001]. **C:** Unpaired context fear expression test. Mice did not generalize fear to the novel context when they were trained in an ‘alternative’ training context and exposed to the ‘standard’ context six hours later. Freezing was higher during the test in the standard training context than in the novel context (p = 0.0275), likely due to some transfer of learning during the ‘explicitly unpaired’ exposure on the acquisition day. There was a significant difference between the ‘standard’ training context and the ‘alternative training context’ (p = 0.0132), as well as the novel context and the ‘alternative training context’ (p < 0.0001). Freezing in the ‘standard’ training context did not differ between training and the expression test (p = 0.1004). Mice that received immediate shock in the ‘standard’ training context and tested in a novel context also displayed low freezing, suggesting that generalization is not strictly due to shock exposure without a representation of context.

#### NMDAR-dependent mechanisms in the ACC are necessary to encode general information about a specific contextual fear experience to support generalization

To further test the idea that associative plasticity in the ACC during fear learning accounts for generalized learning, we utilized the competitive NMDA receptor antagonist, DL-AP5. Male and female mice received DL-AP5 or vehicle immediately before undergoing context fear conditioning. We found no significant difference between treatment groups during acquisition [F (1, 25) = 0.1467, p = 0.7049] (Figure 3B). There were no differences in fear acquisition or expression in the training or novel contexts between male and female mice (Figure S1, S1B; 3-way ANOVA no main effect of sex, F(1, 22) = 2.385, p = 0.14). Therefore, all data was collapsed across sex. During the drug-free fear recall tests, mice that were administered AP5 during training displayed significantly less freezing in the novel context compared to mice administered vehicle control. AP5 had no effect on freezing in the training context (main effect of treatment [F (1, 24) = 9.660, p = 0.0048]; main effect of context [F (1, 24) = 243.4, p < 0.0001]; significant interaction effect [F (1, 24) = 4.986, p = 0.0351]; Sidak’s p = 0.0008) (Figure 3C). We calculated freezing difference scores between the training and novel contexts using the formula [novel context freezing – training context freezing] / [novel context freezing + training context freezing] as a generalization index. AP5-treated mice had a significantly reduced generalization index (i.e., greater freezing difference between the training and novel context) compared to vehicle-treated mice [t (24) = 3.14, p = 0.0044] (Figure 3D), indicating reduced generalization. Given that NMDA receptors are necessary for long-term potentiation and associative learning (*see* Kandel, 2009), these data suggest associative plasticity in the ACC is necessary for mice to encode general contextual elements that promote generalized fear during recall tests in a novel context. Importantly, post-training infusions of AP5 had no effect on freezing in either context (main effect of context only [F (1, 35) = 41.01, p < 0.0001; Tukey’s HSD, p = 0.96) (Figure 3E), consistent with data suggesting that NMDA receptors are only necessary for the induction of LTP and not its maintenance (Collingridge, et al., 1983; Bliss & Collingridge, 1993). It is important to point out that NMDAR blockade did not affect learning to the specific training context, suggesting that the role of NMDAR-dependent plasticity in the ACC is to encode general information that supports memory generalization.

**Figure 3:**
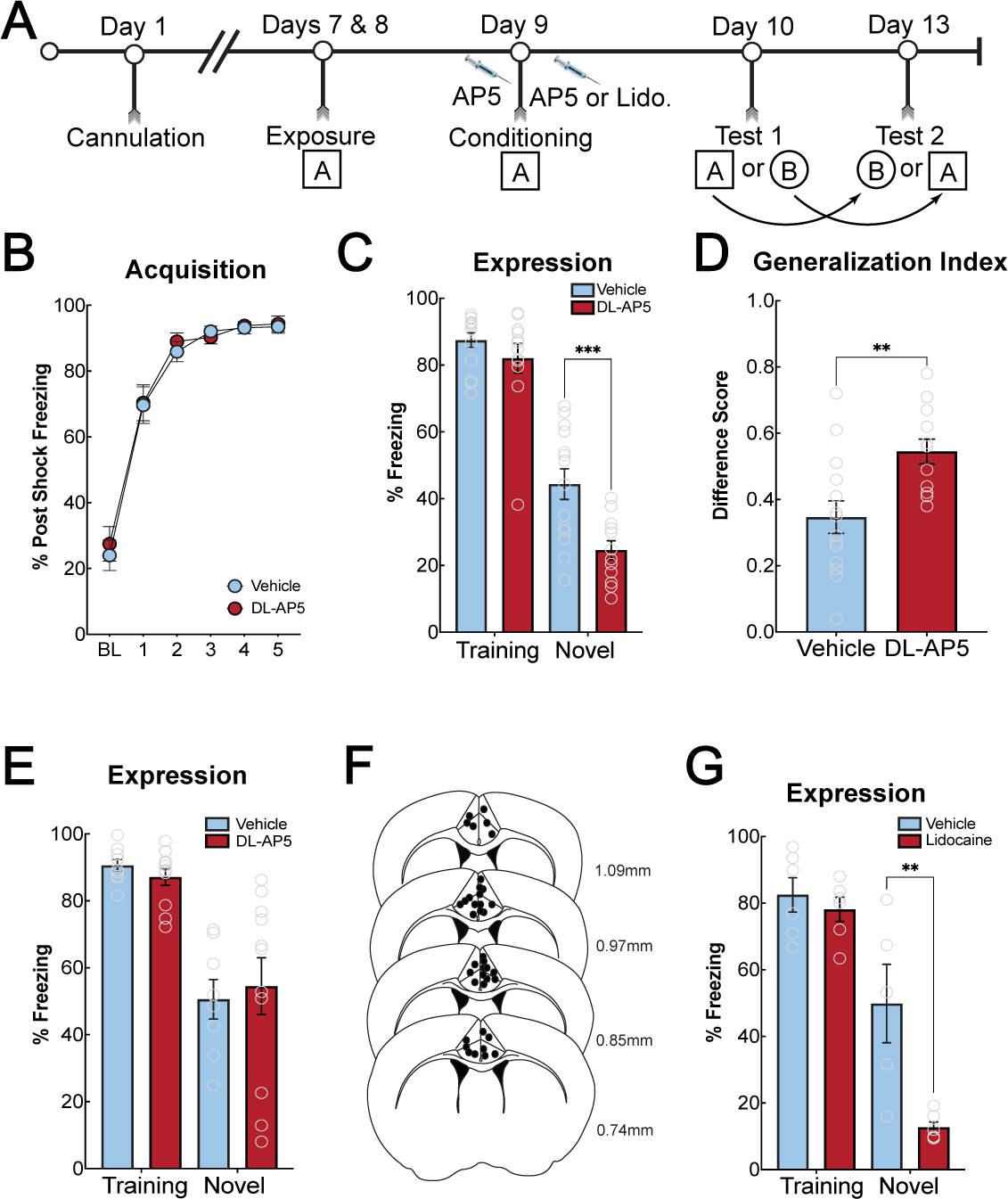
Inactivation of NMDAR in the ACC blocks the acquisition of generalized context fear. **A:** Timeline of behavioral experiments. **B:** Acquisition of context fear conditioning during the experiment in which mice were infused with DL-AP5 before training. There were no differences between DL-AP5 and vehicle-treated mice during acquisition. **C:** NMDAR blockade in the ACC during learning significantly attenuates contextual fear generalization. Mice receiving DL-AP5 during training displayed significantly less freezing to the novel context than vehicle-treated mice (p = 0.0008). Freezing to the training context was not different between the groups.**D:** Generalization index for pre-training DL-AP5. AP5-treated mice had a significantly greater difference score compared to vehicle-treated mice, indicating a greater difference between their training context freezing compared to their novel context freezing [t (24) = 3.14, p = 0.0044].**E:** Post-training NMDAR blockade in the ACC does not affect context fear generalization or context-specific fear. There were no significant differences in freezing between AP5- and vehicle-treated mice in the novel (p = 0.9629) or training context (p = 0.9719). **F:** Schematic of cannula placements in the ACC. Placements for pre-training and post-training DL-AP5 experiments are combined into one schematic. **G:** Post-training inactivation of the ACC with lidocaine blocked context fear generalization but had no effect on context-specific fear. Mice infused with lidocaine immediately after training showed a significant reduction in freezing to the novel context compared to vehicle-treated mice (p = 0.0025). There were no differences in freezing in the training context (p = 0.9531).

#### The ACC consolidates general information about a specific contextual fear experience to facilitate memory generalization

Given that NMDAR-dependent plasticity in the ACC is required to acquire a generalized fear memory, we next wanted to determine if the ACC was involved in consolidating newly acquired generalized memories. To accomplish this, we used a post-training manipulation, which is typically thought to influence the consolidation of memory (Power, et al., 2003; LaLumiere, et al., 2005; Malin, et al., 2007). Mice were trained in the context fear task as above and received lidocaine immediately after training to reversibly inactivate the ACC. Twenty-four hours after training, mice were tested in the training or novel contexts. Post-training lidocaine infusions in the ACC eliminated generalized fear to the novel context but left specific fear to the training context intact (main effect of context [F (1, 19) = 64.52, p < 0.0001], main effect of treatment [F (1, 19) = 11.58, p = 0.0030], significant interaction [F (1, 19) = 7.206, p =0.0147], Tukey’s HSD, (p = 0.0025) (Figure 3G). These experiments demonstrate that the ACC is critical for encoding and consolidating generalized context memory associated with elevated threat. The ACC, however, is not necessary to acquire or consolidate specific contextual memory of the training context, at least under the training conditions used here.

### Generalized contextual memory formation requires the ACC, but not PL

We were next interested in understanding if the ACC plays a unique role in regulating generalized contextual memory formation or if this process involves multiple prefrontal cortical regions. We investigated the prelimbic cortex (PL; A32) given its role in fear expression (Corcoran & Quirk, 2007; Sierra-Mercado, et al., 2011) as well as regulating discrimination between aversive and non-aversive cues (Stujenski, et al., 2022). Mice received bilateral infusions of inhibitory DREADD virus hM4Di or an EGFP control. Five weeks after surgery, mice underwent contextual fear conditioning using the abovementioned procedure. Thirty minutes prior to training, all mice received an i.p injection of CNO to inactivate the PL. There was no difference in acquisition between the groups (main effect of shock only [F (2.981, 44.71) = 95.18, p < 0.0001], Figure 4D). Mice received counterbalanced tests in the training and novel contexts. We found a significant main effect of context [F (1, 15) = 23.55, p = 0.0002]. However, Sidak’s posthoc analysis did not reveal a significant difference between hM4Di or EGFP for the training (p = 0.9970) or novel (p = 0.9854) contexts (Figure 4E). The generalization index was also not different between the groups [t (15) = 0.1653, p = 0.8709] (Figure 4F). This emphasizes that the ACC is important for regulating generalized contextual memory formation and is not a general function of the mPFC. However, the PL and infralimbic cortex (IL) are implicated in explicit cue generalization (Stujenske, et al., 2022; Pollack, et al., 2018; Likhtik, et al., 2014). This suggests that the PL is important for regulating fear responses based on appropriate contextual cues. In contrast, the ACC may be more involved in context-independent responses, enabling animals to generalize responses across different contexts.

**Figure 4:**
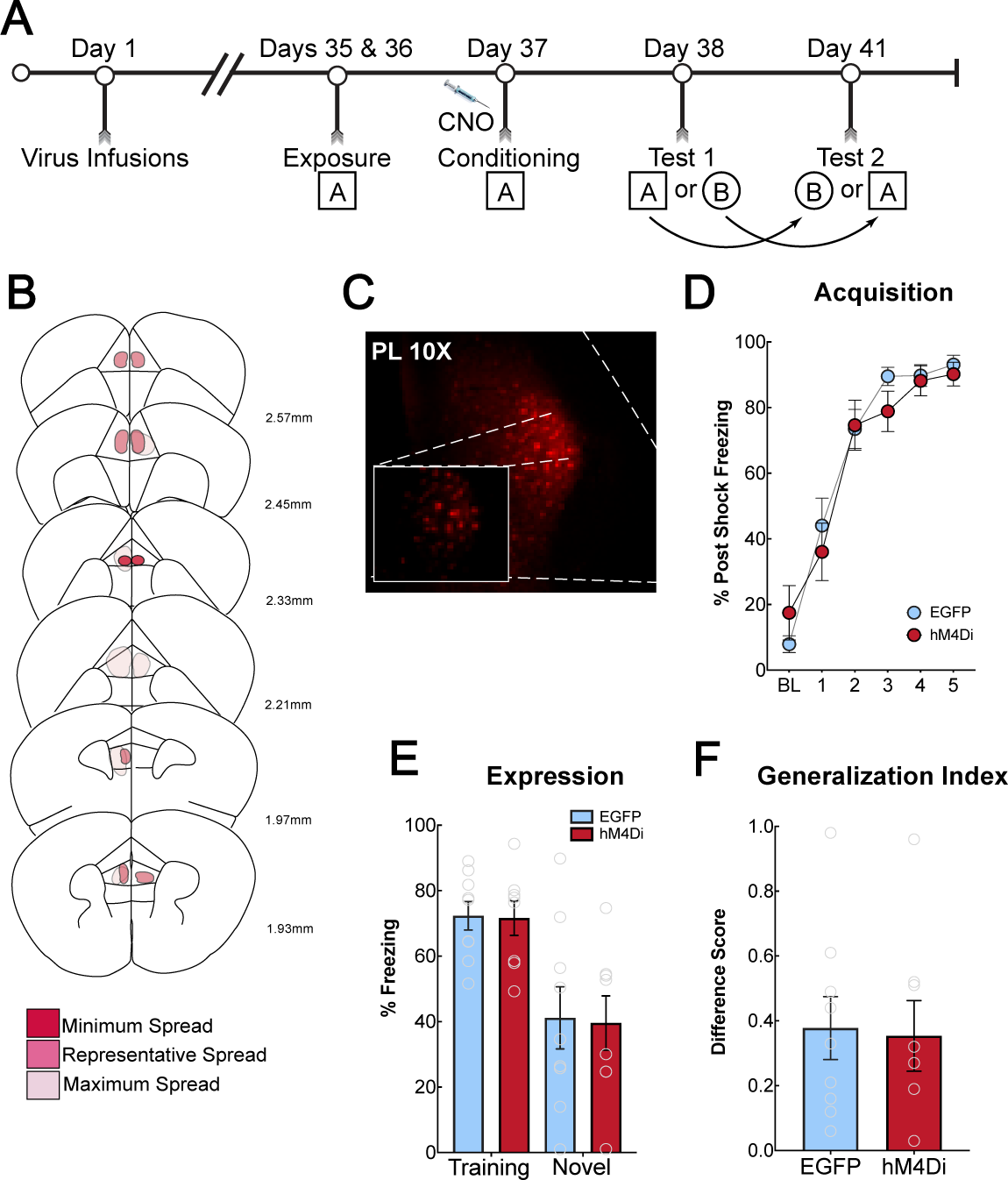
Chemogenetic inactivation of the prelimbic cortex during training does not reduce context fear generalization. **A:** Timeline of behavioral experiment. Mice were infused with an hM4Di-mCherry or mCherry-expressing control AAV and received injections of CNO before context fear conditioning. Mice were tested for fear responses in the training (square, A) and novel (circle, B) contexts in a counterbalanced design. **B:** Schematic representation of hM4Di-mCherry expression in the prelimbic cortex (PL). Dark pink represents minimum virus spread, medium pink represents average, and light pink represents maximum virus spread. **C:** Representative image of PL hM4Di-mCherry expression. **D:** During acquisition, mice expressing hM4Di or mCherry showed no differences in their post-shock freezing. **E:** Mice were tested for fear expression in the training and novel context with 72-hours between tests. There were no differences in freezing in the training (p = 0.9970) or novel context (p = 0.9854) between hM4Di- and mCherry-expressing mice. **F:** Generalization index for PL inactivation. There were no differences in the generalization index between hM4Di-expressing and mCherry-expressing mice.

### BLA projections to the ACC are necessary for encoding general threat information to promote generalized context fear

We next investigated circuit mechanisms underlying how the ACC encodes generalized fear memories. For the ACC to encode and consolidate generalized memories, it would likely need to receive threat information from the amygdala. While additional regions could provide contextual or nociceptive information such as the hippocampus and thalamus, we first chose to investigate the inputs from the basolateral amygdala (BLA), given its critical role in fear acquisition, consolidation, and expression (Alvarez, et al., 2008; Helmstetter, et al., 1994; Maren, et al., 1996; Maren, 1999; Fanselow & LeDoux, 1999; *for review see* Tovote, et al., 2015). We utilized an intersectional chemogenetic approach bilaterally injecting a retrograde cre-expressing virus in the ACC (pAAV-Ef1a -cre (AAVrg)) and a cre-dependent virus encoding hM4Di or mCherry bilaterally in the BLA (pAAV-hSyn-DIO-hM4D(Gi)-mCherry (AAV8) or pAAV-hSyn-DIO-mCherry (AAV8)). Five weeks after surgery, mice underwent contextual fear conditioning. Thirty minutes prior to training, all mice received an i.p injection of CNO. Male and female mice acquired fear similarly (Figure 5F). Mice were tested in the training or novel context in a counterbalanced design with 72 hours in between tests. BLA-to-ACC circuit inactivation eliminated generalization in a novel context, but specific fear to the training context was unaffected ([F (1, 22) = 9.014, p = 0.0066]). We found no differences between males and females; therefore, the data were collapsed and analyzed together (3-way ANOVA; main effect of context [F(1, 20) = 249.4, p < 0.0001]; main effect of virus [F(1, 20) = 7.535, p = 0.0125], no main effect of sex [F(1, 20) = 3.472, p =0.0772], Figure S2). Post-hoc analysis revealed a significant difference between hM4Di and mCherry groups for freezing in the novel context (p = 0.0031) (Figure 5G). hM4Di-expressing mice also had a significantly reduced generalization index (i.e., higher freezing difference score compared to mCherry controls) [t (22) = 2.693, p = 0.0133] (Figure 5H), suggesting reduced generalization. To verify that inactivation of the BLA-to-ACC circuit did not alter locomotor behavior, we assessed the inactivation of this circuit on locomotion using an open field. Mice received i.p. injections of CNO 30 minutes prior to being placed in the open field. There was no significant difference in distance traveled between mCherry and hM4Di expressing mice [t (17) = 0.2473, p = 0.8076] (Figure 5I). We also saw no differences for time spent in the center [t (17) = 0.8396, p = 0.4128], indicating a lack of effect on anxiety-like behavior. These results indicate that the BLA’s inputs to the ACC during learning, are critical in providing threat information that is utilized by the ACC to encode and consolidate generalized contextual fear memories under increased levels of threat.

**Figure 5:**
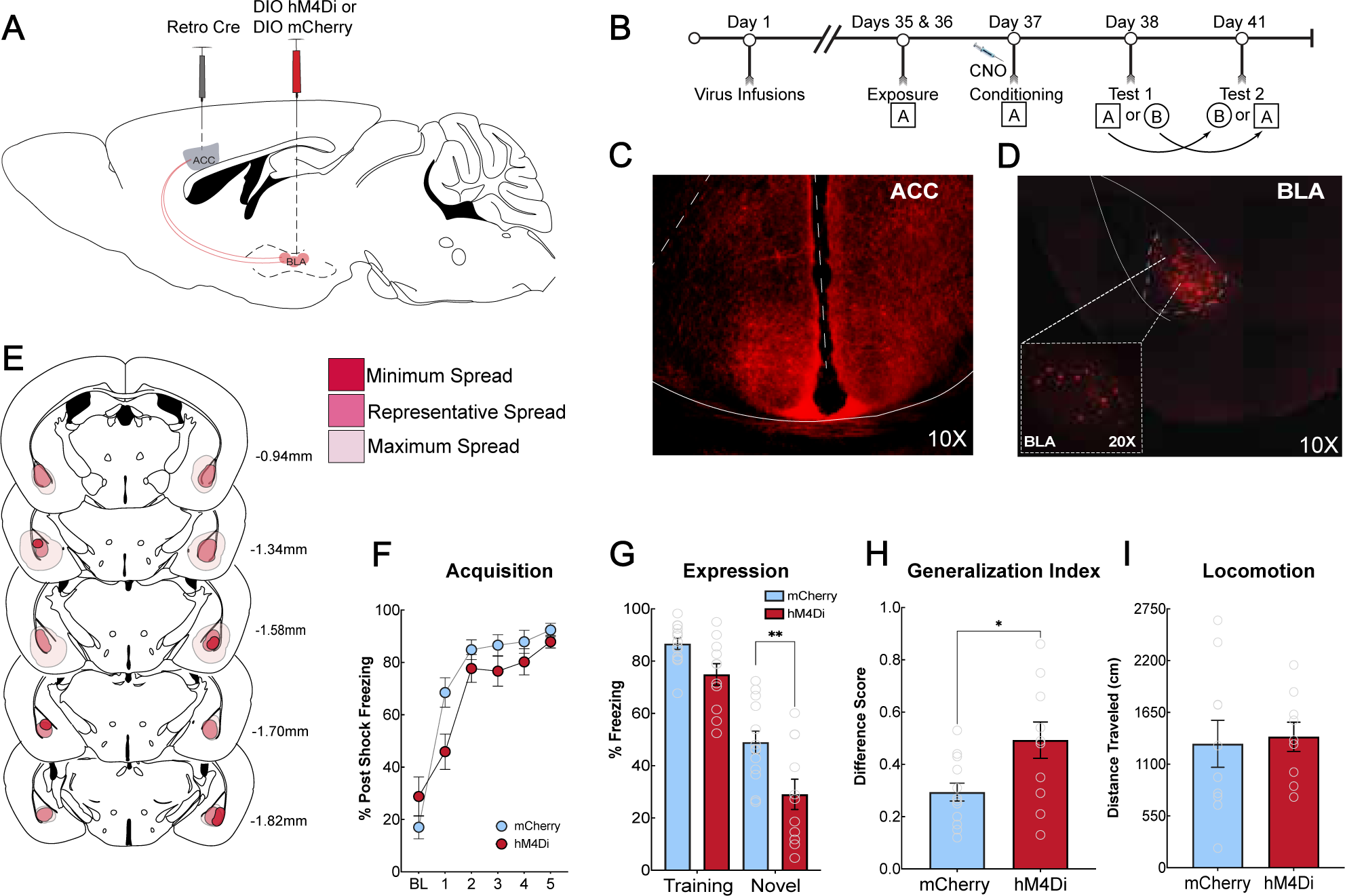
BLA Inputs to the ACC are necessary to encode context fear generalization. **A:** Schematic of viral inactivation strategy. A retrograde AAV expressing cre recombinase was bilaterally injected into the ACC and a cre-dependent AAV expressing hM4Di or mCherry (DIO hM4Di-mCherry or DIO mCherry) was bilaterally injected into the BLA. This enabled specific inactivation of BLA neurons projecting to the ACC. **B:** Timeline of the behavioral experiment. After viral infusions, mice underwent context fear conditioning and were tested for fear responses in the training (square, A) and novel (circle, B) contexts in a counterbalanced design. **C:** Representative image of fiber expression in the ACC. A 10X confocal image was acquired. Image shows hM4Di-mCherry fiber expression in the ACC demonstrating fibers from BLA projection neurons. **D:** Representative image of hM4Di-mCherry expression in the BLA. Images were obtained on a stellaris confocal microscope with a 10x objective (main image). Inset is a 20x image of neurons within the BLA. **E:** Schematic of hM4Di viral spread in the BLA. Dark pink represents minimum virus spread, medium pink represents average, and light pink represents maximum virus spread. **F:** Inactivation of BLA-to-ACC circuit during learning. All mice showed increased post-shock freezing with each shock delivery and there were differences between the groups. **G:** Inactivation of the BLA-to-ACC circuit during context fear learning is necessary for context fear generalization. BLA-to-ACC circuit inactivation eliminated generalization in the novel context, but specific fear to the training context was unaffected (main effect of context [F (1, 22) = 9.014, p = 0.0066] Sidak’s post hoc, p = 0.0031). **H**: Generalization index for hM4Di-expressing and mCherry-expressing control mice. hM4Di-expressing mice had greater freezing difference scores between the training and novel context compared to mCherry-expressing controls [t (22) = 2.693, p = 0.0133]. **I:** Inactivation of BLA-to-ACC circuit did not alter locomotion in the open field. Mice received i.p. injections of CNO (5mg/kg) 30 minutes prior to being placed in the open field. We saw no significant difference in distance traveled between hm4Di- and mCherry-expressing mice [t (17) = 0.2473, p = 0.8076].

### The strength of threat information from the BLA-to-ACC pathway determines the encoding of context fear generalization

We next wanted to test if threat information carried by BLA projections to the ACC could drive encoding of generalization under standard training conditions. First, we ran naïve mice through a standard training protocol consisting of 3 un-signaled foot shocks delivered at 0.6mA, or the strong training protocol used earlier. Twenty-four hours following training, mice were tested in a novel context to assess context generalization. Mice that received strong training (5 shocks, 1.0mA) froze significantly higher compared to mice exposed to the standard training protocol (3 shocks, 0.6mA) [t (16) = 2.508, p = 0.0233] (Figure 6C), indicating that mice trained using the strong training protocol generalized fear to the novel context, whereas the mice trained using the standard protocol did not. Therefore, these training parameters were used to test the sufficiency of the BLA’s inputs to the ACC during context fear learning. Mice were injected with a retrograde virus in the ACC that expressed the chemogenetic excitatory receptor, hM3Dq, or a control virus encoding mCherry (pAAV-hSyn-hM3Dq-mCherry (AAVrg); pAAV-CaMKIIa-EGFP (AAVrg)). Five weeks later, mice were bilaterally cannulated over the BLA to enable infusions of CNO to locally activate ACC-projecting BLA neurons. Following an additional week of recovery, mice were run through context fear conditioning utilizing the standard training protocol. Five minutes prior to training, all mice received infusions of 0.2μL of CNO into the BLA per hemisphere (Ortiz, et al., 2019). There were no differences in fear acquisition between the treatment groups (main effect of shock [F (2.294, 25.24) = 19.61, p < 0.0001]) (Figure 6D). Twenty-four and seventy-two hours after training, mice were tested in the training and novel contexts in a counterbalanced design. Mice expressing hM3Dq froze significantly more in the novel context compared to those expressing EGFP [Sidak’s post-hoc (p = 0.0161)] (Figure 6E). The generalization index also showed the same effect with hM3Dq-expressing mice displaying more generalization compared to EGFP-expressing mice [t (11) = 3.481, p = 0.0051] (Figure 6F). Collectively, these results suggest that inputs from the BLA to the ACC convey threat information, and activation of this input is critical for mice to encode general information regarding a threatening experience. This pathway is likely part of a larger circuit that is recruited to enable animals to generalize responses to novel environments, which in turn allows animals exposed to highly threatening experiences to quickly assess similar situations for the likelihood of threat and respond with appropriate defensive behavior.

**Figure 6:**
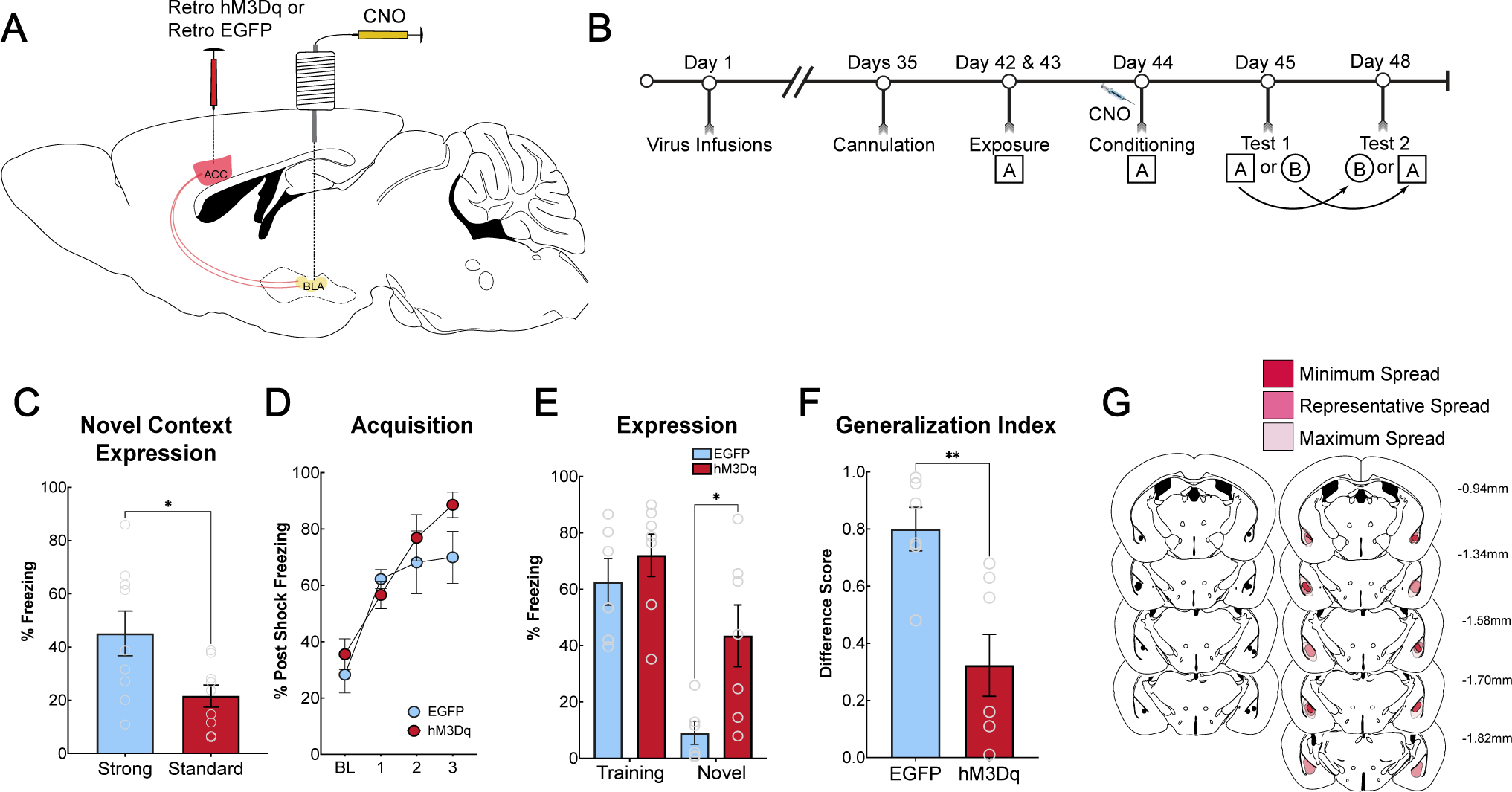
BLA inputs to the ACC drive context fear generalization under training conditions that normally do not support generalization. **A:** Schematic of circuit activation strategy. A retrograde AAV expressing hM3Dq-mCherry or EGFP was bilaterally injected into the ACC. Five weeks later, mice were bilaterally cannulated over the BLA to enable local infusions of CNO in the BLA. One week after cannulations, mice underwent fear conditioning. Infusions of CNO via guide cannula enabled direct activation of BLA neurons that project to the ACC. **B:** Timeline of behavioral experiments. Mice received a bilateral infusion of CNO into the BLA immediately before undergoing context fear training. They were then tested in the training and novel context in a counterbalanced design. **C:** Mice were trained with either a three-shock, 0.6mA protocol (standard training) or a 5 shock, 1.0mA (strong training). There was a significant difference in freezing to the novel context (p = 0.0233). Mice trained with standard fear conditioning displayed less freezing to the novel context than mice with strong training. Mice that underwent the strong training procedure displayed the expected generalization to the novel context. **D:** When the BLA-ACC circuit was activated using chemogenetics during training, both groups of mice acquired context fear as expected, with no differences between treatment groups (main effect of shock [F (2.294, 25.24) = 19.61, p < 0.0001]). **E:** During a fear expression test, hM3Dq-expressing mice froze significantly more in the novel context compared to EGFP-expressing mice (main effect of context [F (1, 11) = 49.89, p < 0.0001], Sidak’s post-hoc (p = 0.0161)). **F:** The generalization index indicated a greater difference in freezing between the training context and novel context in EGFP-expressing mice compared to hM3Dq-expressing mice [t (11) = 3.481, p = 0.0051]. **G:** Schematic of cannula placements in the BLA and hM3Dq-mCherry in the BLA.

## Discussion

In rodent fear literature, the ACC has been best characterized for its role in remote contextual fear (Frankland, et al., 2004; Frankland, et al., 2006; Vetere, et al., 2011; Goshen, et al., 2011; Einarsson, et al., 2012; 2015). However, we and others have consistently shown a role for the ACC in regulating generalized fear expression both at remote and recent testing periods after conditioning (Cullen, et al., 2015; Einarsson, et al., 2015; Adkins, et al., 2019; Ortiz, et al., 2019). Clinical investigations have found a role for both the ACC and the BLA in anxiety and stress-related disorders (Shin, et al., 2009; Asami, et al., 2008; Liberzon, et al., 2015; Admon, et al., 2013). The ACC within these studies is analogous to the human midcingulate cortex, often referred to as just anterior cingulate cortex within clinical studies (van Heukelum, et al., 2020). Additionally, hyperfunction of the ACC and BLA has been shown to be a predisposition for developing PTSD (Admon, et al., 2013), and overgeneralization of fear is a core symptom of this disorder (Dunsmoor, et al., 2011; Dymond, et al., 2015). Our findings show that the ACC is recruited during training, where Arc activity was highest compared to our control groups (Figure 1C). Next, we identified that NMDAR-dependent plasticity is necessary, which we hypothesize is evoked by inputs from the BLA, to allow for generalized fear expression 24 hours after fear learning. Combined with the post-training inactivation results, our data suggests that the ACC consolidates a general representation of the training experience to guide fear in novel environments (Figure 3). Finally, we found that BLA-to-ACC inputs are both necessary and sufficient for encoding generalized representations (Figures 5 and 6).

To decipher the role of the ACC in processing generalized information, we first investigated its activity following strong training, which produces generalized fear responses (Ortiz, et al., 2019). The ACC has been implicated in processing pain information (Iqbal, et al., 2023; Bliss, et al., 2016; Lee, et al, 2022; Smith, et al., 2021; Johansen, et al., 2001), therefore we utilized an immediate shock procedure, using the same shock parameters as our strong training protocol (Wiltgen, et al., 2001; Fanselow, 1990), to demonstrate that shock alone is not sufficient to drive ACC activity and fear generalization. When we measured ACC Arc protein expression, we saw that the immediate shock group had significantly fewer Arc-expressing cells compared to the strong training group (Figure 1C), however, mice that underwent immediate shock showed significantly more Arc expression than home cage controls in the ACC as would be expected because of the ACC’s role in pain processing. Mice exposed to immediate shock did not generalize fear to the novel context, emphasizing that strong shock exposure alone does not produce generalization. Using an unpaired training procedure in which mice received shocks and context exposure separately, we further showed that nociceptive pain processing alone is not sufficient to drive context generalization (Figure 2C).

Next, we utilized pre-training infusions of DL-AP5, an NMDA receptor antagonist to target memory encoding. Long-term potentiation is thought to be the basis of consolidation of memories (*for review, see* Kandel, 2009). NMDA receptors are generally necessary for the induction of plasticity but not for its maintenance across time (Bliss & Collingridge, 1993; Collingridge, et al., 1983). Therefore, using pre-training inactivation of NMDA receptors in the ACC targets the induction period for LTP. In contrast, post-training manipulations would fail to inactivate the receptors when their activity is necessary. We found that inactivation of NMDA receptors in the ACC prior to learning significantly reduced fear in the novel context but left fear in the training context intact (Figure 3C). We utilized a post-training NMDA receptor manipulation and found no effect on generalized fear (Figure 3E). These data suggest that the mechanisms within the ACC that drive generalized context fear are mediated through NMDAR-dependent synaptic plasticity. Furthermore, immediate post-training inactivation of the ACC significantly attenuated context fear generalization, suggesting that cellular activity within the ACC following learning is critical for generalization (Figure 3G). We next verified that our results are restricted to the ACC, and not the PL, given its role in fear expression (Corcoran & Quirk, 2007; Sierra-Mercado, et al., 2011) as well as discriminating between aversive and non-aversive cues (Stujenski, et al., 2022). We used chemogenetics to inactivate the PL prior to strong training and found no effect on the acquisition (Figure 4D) or during the test in the training or novel context (Figure 4E). Overall, these data demonstrate that the ACC is recruited to encode and consolidate generalized fear memories, which are not reliant on the PL.

Whereas we consistently observe a selective role for the ACC in regulating generalized context fear, several previous studies have demonstrated that the ACC is important for consolidating or expressing fear in the training context. For example, inactivation of the ACC disrupts the retrieval of remote context fear memory (Frankland et al., 2004; Goshen et al., 2011). In addition, the administration of Ro25 or NR2B subunit siRNA into the ACC before learning decreases fear expression in the training context; the administration of anisomycin also produced the same effects (Einarsson et al., 2012; Zhao et al., 2005), suggesting an important role of NMDAR-dependent plasticity and cellular consolidation within the ACC for contextual memory. Here, we show that the ACC is selective in encoding and consolidating generalized contextual memory but not memory for the training context. It is difficult to reconcile the differences among these studies and why some have found a selective role for the ACC in generalized context memory (e.g., Cullen et al., 20014, Einarsson et al., 2015, Ortiz et al., 2019), and others have found it to regulate consolidation and expression of specific context memory (e.g., Einarsson et al., 2012, Zhao et al., 2005, Frankland et al., 2004). One difference could be our use of context pre-exposures in this and our previous studies. However, while several studies have not used context pre-exposures, Einarsson et al., 2012 did use context pre-exposure and found the ACC to be important for the consolidation and expression of specific context fear. It has been suggested that NMDAR-dependent mechanisms in the ACC are recruited for newly acquired information or if prior learning does not match the new experience (Finnie, et al., 2018; Xia & Frankland, 2018). Thus, our results could be due to the recruitment of NMDAR-dependent mechanisms in the ACC because the previously acquired pre-exposures no longer match the training due to the addition of the shocks. However, Einarsson et al., 2012 also used a context pre-exposure and found the ACC to regulate memory for the training context and reconsolidation of that memory. Additional specific methodological differences could explain some of the discrepancies with some, but not all, of the studies; our manipulations before or after training were performed without anesthetizing mice, as in Frankland et al., (2004). In another case, the authors performed tone-dependent fear training with context as background and used multiple recall tests in the same context (Goshen et al., 2011). Here, we used unsignaled shocks to train specifically for contextual fear, and mice were only tested in a single context once. Finally, Zhou et al., (2005) used single footshock conditioning during training, whereas we used 5 unsignaled footshocks. These discrepancies suggest that the ACC involvement in specific versus generalized fear responses could be related to the strength and type of training.

The BLA is known for its role in both the acquisition and expression of fear (Alvarez, et al., 2008; Helmstetter, et al., 1994; Maren, et al., 1996; Maren, et al., 1999; Huff & Rudy, 2004; Fanselow & LeDoux, 1999; *for review see* Tovote, et al., 2015), making it a critical node for both receiving and sending information about a threatening experience. The BLA and ACC share reciprocal connections (Hoover & Vertes, 2007; Hintiryan, et al., 2021), and we have previously identified that ACC efferent projections to the BLA regulate generalized context fear expression (Ortiz, et al., 2019). Here, we investigated the role of the ACC itself and ascending BLA projections to the ACC during context fear learning. We utilized an intersectional chemogenetic approach to inactivate BLA inputs to the ACC. Before DREADD experiments, we trained surgical naive mice using contextual fear conditioning and administered CNO before learning. We found that CNO alone, without DREADD virus expression, did not affect fear learning (Figure S3B) or fear recall in both the training and novel context and did not affect locomotion (Figure S3C). Next, we administered CNO prior to learning. We found a significant reduction of freezing in the novel context but not the training context (Figure 5G), suggesting that BLA-to-ACC projections are critical in promoting the encoding of a generalized fear memory. In contrast, activating BLA inputs to the ACC induced generalization to the novel context (Figure 6E). Taken together these data indicate that the BLA-to-ACC circuit is both necessary and sufficient for encoding context fear generalization. The amygdala is thought to assign positive and negative valence (Beyeler, et al., 2018; O’Neill, et al., 2018) to events that evoke strong emotional responses, such as fear (*for review see* Tye, 2018 and Janak & Tye, 2015). In contrast, the ACC is implicated in regulating action selection (Akam, et al., 2021; Rolls, 2019; Hadland, et al., 2020) and decision-making (Kennerley, et al., 2006). Therefore, inputs from the BLA to the ACC likely convey threat information associated with aversive stimuli that engage the ACC to initiate adaptive behavioral strategies in uncertain situations, such as placement into a novel context.

Our procedure utilized two days of pre-exposure, which allows the animals to form a strong representation of the training context (Biedenkapp, J.C., & Rudy, J.W., 2007; Fanselow, et al., 1990). Therefore, the level of fear we observe in the novel context is most likely due to the strong training parameters and similarity of contextual features that produce generalization, not a poor representation of the training context, as would be expected if the mice were not pre-exposed for several minutes. The cortex processes contextual information (Zelikowsky, et al., 2013; Coelho, et al., 2018; Heroux, et al., 2017; Maren & Fanselow, 1997), and in the absence of a precise hippocampal memory, it has been suggested that an impoverished representation is formed (Ramanathan, et al., 2018). Therefore, the ACC could be at least one region where an impoverished context representation can be encoded after high threat levels that would facilitate generalized responding across novel but similar contexts.

BLA-to-ACC inputs have been shown to become active following pain, as well as being linked to pain-induced depressive-like behaviors (Becker et al., 2023). Therefore, plasticity in the ACC may be partially evoked by pain during context fear learning. However, the Arc expression suggests that the ACC is more active following context fear conditioning than immediate shock, which represents the activity of the shock alone without an association (Figure 1). In addition, with standard conditioning, mice fail to generalize fear the following day (Figure 6). However, in the presence of strong training, BLA inputs to the ACC drive threat-related plasticity. It is important to note that thalamic subregions send projections to the ACC (Hoover & Vertes, 2007), and it has recently been identified that medial dorsal-to-ACC projections evoke pain-related avoidance (Meda, et al., 2019) as well as alterations in synaptic plasticity on anterior thalamic projections to the ACC following neuropathic pain (Hogrefe, et al., 2022). Therefore, the thalamus could mediate pain-induced plasticity, which is strengthened by recruiting the BLA inputs to the ACC to allow NMDAR-dependent encoding.

In sum, our findings suggest that the ACC may be an integration site for encoding threat information from the amygdala to facilitate generalized behavioral responses. Given that our tests involve shifts in context, we would expect a role for the ventral hippocampus in this process (Ortiz, et al., 2019; Cullen, et al., 2015; Besnard, et al., 2020). However, an interaction between the ventral hippocampus and ACC has not yet been demonstrated in intensity-dependent generalization. The ACC, BLA, and vHPC likely act as a coordinated circuit to contribute to learning and generalizing within the ACC, enabling animals to predict threatening environments and improve survival. Overall, the ACC might integrate information received from several areas to promote flexible behavioral responses following learning. These responses, depending on the context, could be a product of schema or categorical learning, both of which are known to involve the cortex (Freedman, et al., 2001; Tse, et al., 2011; Wang, et al., 2012), which enable animals to respond adaptively to changing environments after highly salient events.

## Authorship Contributions

**CJV** designed research, conducted research, acquired funding, analyzed the data, wrote the manuscript, revised, and edited the manuscript. **RAA** conducted research. **SOV** conducted research. **DDM** designed research and acquired funding. **AMJ** designed research, acquired funding, and revised and edited the manuscript.

## Acknowledgements

We thank the animal care staff at the University of South Carolina School of Medicine.

## Supplemental Materials

**Table 1: Figures 1, 2, 3, & 4 statistical summary.**

**Table 2: Figures 5 & 6 statistical summary. Supplemental Figure Legends:**

### Supplemental Figure Legends

**Supplemental Figure 1:**
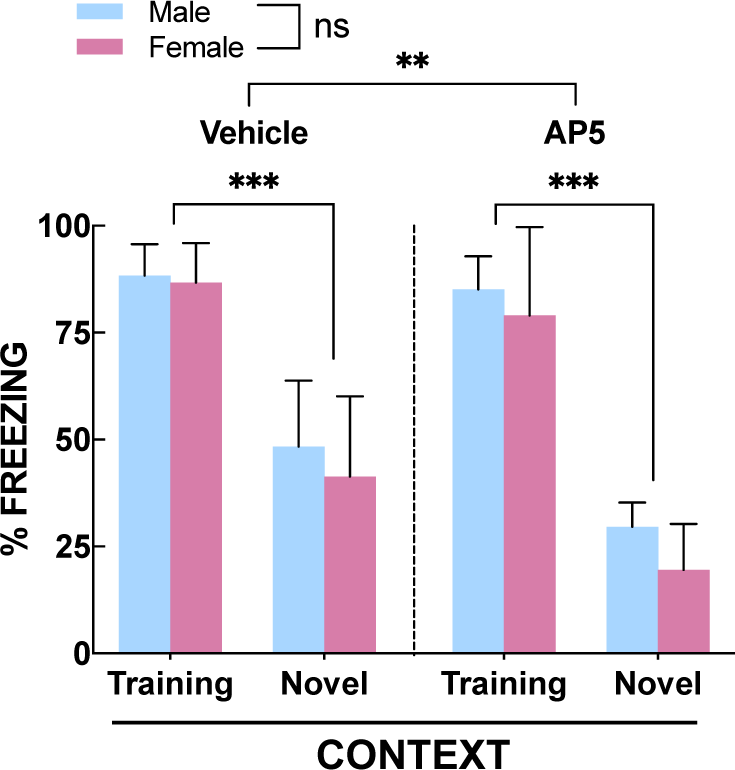
No Sex Differences in NMDAR Inactivation on Fear Generalization. Fear expression test in male and female mice. A 3-way ANOVA (Context x Treatment x Sex) indicated there was a main effect of context (F(1, 22) = 224.4; P<0.0001), a main effect of treatment (AP5 v. vehicle) (F (1, 22) = 10.26; P=0.0041) and a context x treatment interaction (F(1, 22) = 4.878; P=0.0379). There was no main effect of sex (F(1, 22) = 2.385; P=0.1368) and no additional significant interactions.

**Supplemental Figure 2:**
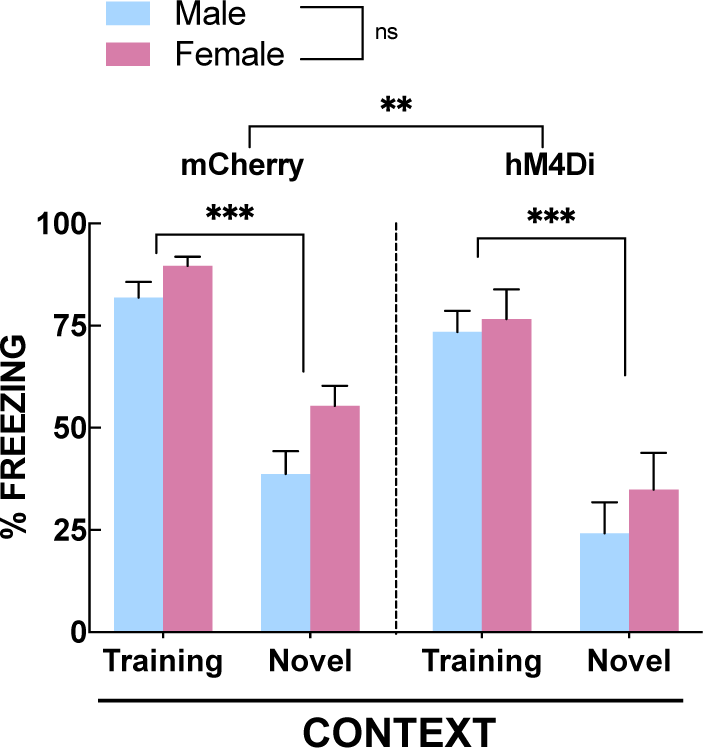
No Sex Differences in BLA-to-ACC Circuit Inactivation on Fear Generalization. A: Fear expression test in male and female mice. A 3-way ANOVA (Context x Treatment x Sex) indicated there was a main effect of context [F(1, 20) = 249.4, p < 0.0001]; a main effect of virus [F(1, 20) = 7.535, p = 0.0125], but no main effect of sex [F(1, 20) = 3.472, p =0.0772].

**Supplemental Figure 3:**
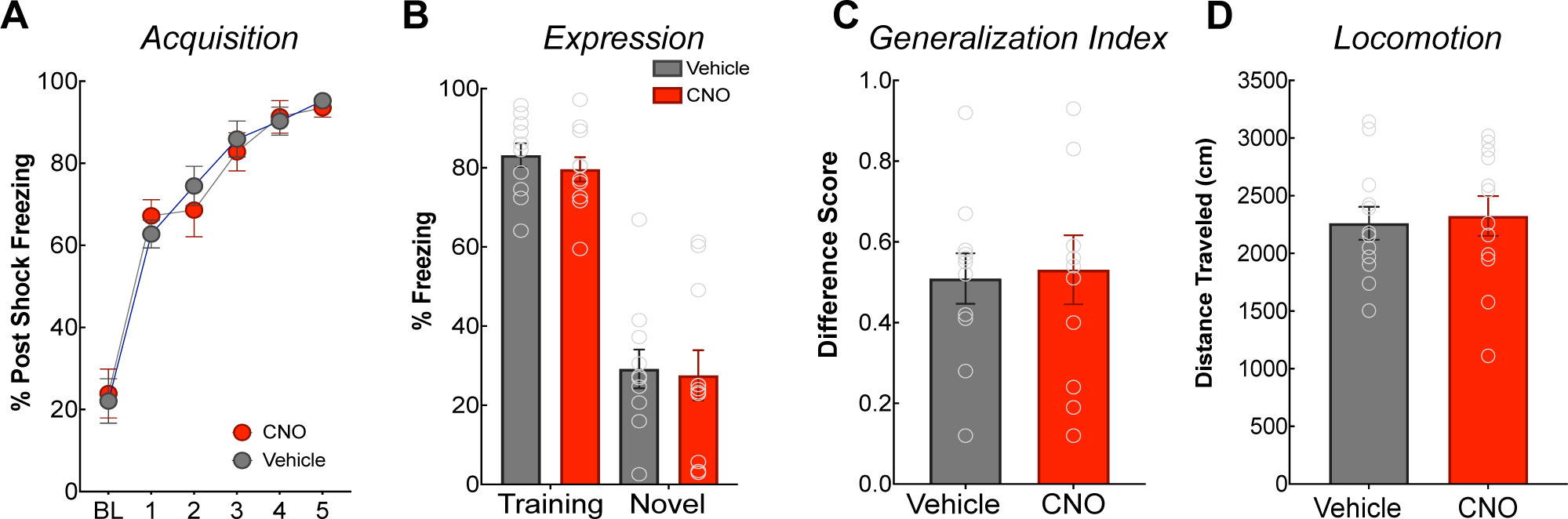
CNO Itself does not affect Fear Learning, Recall, Generalization, or Locomotion. Mice were injected i.p. with CNO (5 mg/kg) or vehicle (0.9% sterile saline) 30 minutes before fear conditioning. Mice were fear conditioned with 5 unsignaled footshocks (1 mA), idenditcal to the procedure used for experiments described in the manuscript. **A:** There was no difference between CNO- and vehicle-treated mice in fear acquisition. **B:** Fear expression test in the training context or the novel context. There was no difference in freezing in the training (p = 0.9598) or novel (p = 0.8232) context between CNO- and vehicle-treated mice. **C:** The generalization index was not different between CNO- and vehicle-treated mice [t (20) = 0.2064, p = 0.8386]. **D:** Mice were injected with CNO i.p. 30 minutes before being placed in an open field and their locomotion was recorded for 10 mintes. There was no difference between CNO- and vehicle-treated mice in distance traveled [t (22) = 0.2843, p = 0.7789].

